# Correlation of vaccine-elicited antibody levels and neutralizing activities against SARS-CoV-2 and its variants

**DOI:** 10.1101/2021.05.31.445871

**Authors:** Jinbiao Liu, Brittany H Bodnar, Xu Wang, Peng Wang, Fengzhen Meng, Adil I Khan, A Sami Saribas, Nigam H Padhiar, Elizabeth McCluskey, Sahil Shah, Jin Jun Luo, Wen-Hui Hu, Wen-Zhe Ho

## Abstract

Both Pfizer-BNT162b2 and Moderna-mRNA-1273 vaccines can elicit an effective immune response against SARS-CoV-2 infection. However, the elicited serum antibody levels vary substantially and longitudinally decrease after vaccination. We examined the correlation of vaccination-induced IgG levels and neutralization titers against newly emerged variants remains and demonstrate a significant reduction of neutralization activities against the variants (B.1.1.7, B.1.525, and B.1.351) in Pfizer or Moderna vaccined sera. There was a significant and positive correlation between serum IgG levels and ID_50_ titers for not only SARS-CoV-2 WT but also the variants. These findings indicate that a high level of anti-spike IgG may offer better protection against infection from SARS-CoV-2 and its variants. Therefore, it is necessary to longitudinally monitor specific serum IgG level for evaluating the protective efficacy of the vaccines against SARS-CoV-2 and its new variants.

---

The COVID-19 vaccines (Pfizer-BNT162b2 and Moderna-mRNA-1273) can elicit an effective immune response against severe acute respiratory syndrome coronavirus 2 (SARS-CoV-2) infection ^1,2^. However, titers of elicited serum antibody and neutralizing activities against SARS-CoV-2 vary considerably among vaccinated individuals and decline after vaccination ^3,4^. Additionally, the protective efficacy of the vaccines against newly emerged variants remains to be elucidated. Therefore, it is of importance to understand the correlation between levels of vaccine-induced antibody and neutralizing activity against SARS-CoV-2, including the variants.

## Methods

This study was approved by Temple University IRB (IRB #28021) and the informed consent forms were signed by all study subjects. Sera samples were obtained from 30 Pfizer vaccinated subjects (22-68 days after 2^nd^ dose) and 19 Moderna vaccinated subjects (24-49 days after 2^nd^ dose) (**Supplemental content Table S1**). Peripheral blood samples were collected three weeks to two months after the second dose of vaccine. The serum titers of specific IgG antibody to SARS-CoV-2 spike S1 by an enzyme-linked immunosorbent assay (ELISA).

Neutralization assays were performed with recombinant vesicular stomatitis virus (rVSV)-based pseudoviruses bearing SARS-CoV-2 spike proteins from the original Wuhan-1 reference isolate (Wild Type, WT) and the variants (D614G, UK-B.1.1.7, UK-B.1.525, and SA-B.1.351) (**Supplemental content Table S2**). Serum neutralizing titers (50% inhibitory dilution, ID_50_) of all vaccinated subjects were determined using four-parameter logistic curve. Geometric mean titers (GMTs) were calculated with 95% CI. Wilcoxon matched-pairs signed rank test was used for two-group analysis. Pearson’s correlation coefficients were calculated. *P* values less than 0.05 were statistically significant.

## Results

There was a broad distribution of IgG levels among the vaccinated subjects, ranging from 11,455 ng/mL (lowest level) to 167,989 ng/ml (highest level) with geometric mean of 61,203 ng/mL for Pfizer group and 20,146 ng/ml to 170,270 ng/mL with geometric mean of 92,435 ng/ml for Moderna group (**Supplemental content Fig. S1A**). The differences in geometric mean (61,203 vs. 92,435) between two groups were statistically insignificant.

The neutralizing infectivity of the pseudoviruses was evaluated in sera at dilutions ranging from 1:50 to 1:36,450. We showed that sera from all vaccinated subjects had neutralizing activities and there was no statistical difference in serum neutralizing activity (ID_50_) against SARS-CoV-2 WT between two groups (**Supplemental content Fig. S1B**), although the ID_50_ varied substantially within each group. The Pfizer group ID_50_ ranged from 732 to 30,021 with a geometric mean titer (GMT) of 6,739, and the Moderna group ID_50_ ranged from 2,426 to 26,667 with GMT of 9,670 (**Fig. S1B**).

Although sera from all vaccinated subjects could neutralize the pseudoviruses bearing spike proteins of variants, neutralizing titers were lower when compared to SARS-CoV-2 WT. In Pfizer-vaccinated sera, there was a significant decrease of GMTs for D614G (4,649, −1.45 fold), B.1.1.7 (3,058, −2.2 fold), B.1.525 (1,658, −4.06 fold), and B.1.351 (644, −10.46 fold), respectively (**Fig. 1A**). In Moderna-vaccinated sera, although there was little difference between GMT (9,670) of SARS-CoV-2 WT and that (10,771) of D614G, there was a significant reduction of GMTs for B.1.1.7 (6,525, −1.48 fold), B.1.525 (2,869, −3.37 fold), and B.1.351 (1,279, −7.56 fold), respectively (**Fig. 1B**). Among the variants studied, B.1.351 appeared to be the most resistant to the neutralization by sera from either Pfizer (reduction of 10.46-fold) or Moderna (reduction of 7.56-fold) groups. This finding is consistent with and supported by recent reports ^5,6^. Despite an overall decline in neutralizing titers (GMTs) against the variants, sera at low dilution (1:50) could neutralize 99% of both SARS-CoV-2 WT pseudovirus and the variants (D614G, B.1.1.7, B.1.525, and B.1.351) (**Supplementary Appendix Fig. S2 and S3**). The linear regression analysis showed a significant and positive correlation between serum IgG levels and neutralizing activities (ID_50_) against SARS-CoV-2 WT (R^2^ = 0.2816, *P* < 0.0001) or variants: D614G (R^2^=0.3258, *P*<0.0001), B.1.1.7 (R^2^ =0.2864, *P*<0.0001), B.1.525 (R^2^ =0.2903, *P*<0.0001), and B.1.351 (R^2^ =0.2310, *P*=0.0005), respectively (**Fig. 1C**).

**Figure 1.**
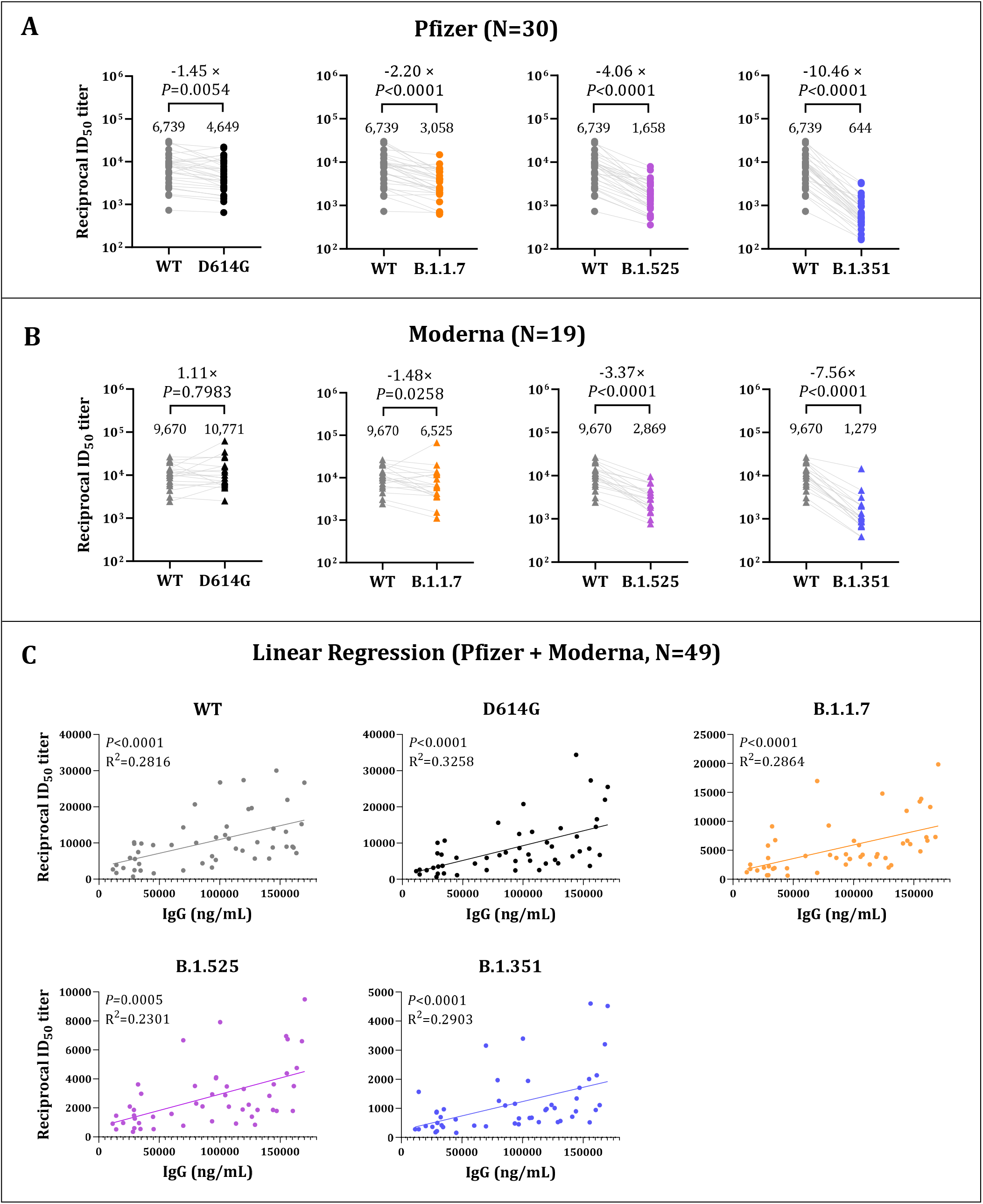
Neutralization of SARS-CoV-2 Pseudoviruses in Sera and its Correlation with Vaccine-elicited IgG Levels. Sera obtained from either Pfzier or Moderna vaccinated subjects were collected three weeks to two months after the second dose vaccine. Neutralizing activity was measured in an assay with recombinant vesicular stomatitis virus (rVSV)-based pseudovirus bearing spike proteins of SARS-CoV-2 WT or the full-set variants. Panels A and B show the reciprocal neutralizing titers at a 50% inhibitory dilution (ID_50_). The lines connect the WT and variant neutralizing titers in matched samples. Fold changes in the reciprocal serum ID_50_ in vaccinated sera against the D614G, B.1.1.7, B.1.525, and B.1.351 variants, as compared with WT, are shown above the *P* value. The dots in Panel A indicate the sera ID_50_ titers of Pfizer vaccinated subjects; the triangles in Panel B indicate the sera ID_50_ titers of Moderna vaccinated subjects. The grey, black, orange, purple and blue symbols represent the ID_50_ titer of the WT, D614G, B.1.1.7, B.1.525, and B.1.351 variant, respectively. The numbers over the dot of each group are the geometric mean titers (GMTs). Panel C shows correlation of the neutralizing titers ID_50_ (abscissa) and anti-SARS-CoV-2 spike S1 IgG levels (ordinate) of sera from vaccinated subjects. (Pfizer, N=30; Moderna, N=19). In Panel A and Panel B, Wilcoxon matched-pairs signed rank test was used for two-group analysis. In Panel C, linear regression analysis was performed using GraphPad Prism 9.1.1. software. Pearson’s correlation coefficients were calculated. Simple linear regression (solid line) is shown. R^2^ = goodness of fit. *P* values less than 0.05 are statistically significant.

## Discussion

This study demonstrated that all study participants vaccinated with either Pfizer or Moderna vaccine were able to produce effective antibodies against spike proteins of both SARS-CoV-2 WT and the variants, although levels of the elicited IgG (specific for SARS-CoV-2 spike protein S1) and the neutralizing titers (ID_50_) varied substantially. There was a several-fold reduction in GMTs of ID_50_ against the variants (UK-B.1.1.7, UK-B.1.525, and SA-B.1.351) in sera, as compared to those against SARS-CoV-2 WT. However, sera at the low dilution were equally effective in neutralizing both SARS-CoV-2 and the variants. More importantly, we demonstrated that among all vaccinated subjects, there was an overall positive correlation between serum IgG levels and ID_50_ titers for not only SARS-CoV-2 WT but also the variants. This finding suggests that a high level of anti-spike IgG may offer better protection against infection from SARS-CoV-2 and its variants. Therefore, it is necessary to longitudinally monitor specific serum IgG level for evaluating the protective efficacy of vaccines against SARS-CoV-2 and its new variants.

Limitations of the study include the small sample size, with possible lack of representativeness, and the lack of live SARS-CoV-2 neutralization assays.

## Supporting information

Supplemental online content

## Author Contributions

Drs Ho, Hu and Luo had full access to all of the data in the study and takes responsibility for the integrity of the data and the accuracy of the data analysis.

Concept and design: Liu, Ho, Hu, Luo.

Acquisition, analysis, or interpretation of data: Liu, Bodnar, Wang, Wang, Meng, Khan, Padhiar, McCluskey. Drafting of the manuscript: Liu, Ho.

Critical revision of the manuscript for important intellectual content: All authors.

Statistical analysis: Liu, Ho.

Administrative, technical, or material support: Wang, Hu, Luo, Ho.

Supervision: Hu, Luo, Ho.

## Additional Contributions

We thank all the voluntary vaccinated subjects for providing clinical samples; members of Neurology Clinic in Temple Hospital for collecting blood samples; Guangxiang Luo, Ph.D. (University of Alabama), for providing Hela/ACE2-11 cells. None of these contributors received any compensation for their help in carrying out the study.

