## Supplemental online content for "Correlation of vaccine-elicited antibody levels and neutralizing activities against SARS-CoV-2 and its variants"

Materials and Methods
eReferences

This supplemental material has been provided by the authors to give readers additional information about their work.

### Materials and Methods

#### Ethical Approval

The study was approved by the Institutional Review Board of Temple University (IRB number: #28021). All the study subjects signed written informed consent.

#### Subjects Enrolled and Human Samples

All the enrolled subjects were given two-doses of either Pfizer or Moderna vaccine on a prime-boost schedule. Peripheral blood samples were collected three weeks to two months after the second dose of vaccine. The sera neutralizing activity against wild type and variants of SARS-CoV-2 was measured.

#### Recombinant VSV-based Pseudovirus Neutralization

A codon-optimized spike protein of the original Wuhan isolate (WT) was cloned into pCAGGS vector. To enhance the packaging efficiency of pseudovirus, the 18 aa at C-terminal of spike was deleted. All the mutations were introduced into the spike gene using one or two steps of NEBuilder HiFi cloning. The plasmids were sequenced to ensure that only the intended mutations were present. To generate recombinant vesicular stomatitis virus (rVSV)-*firefly*-luciferase pseudovirus bearing spike of SARS-CoV-2 S-WT and variants, BHK-21/WI-2 cells (Kerafast, EH1011) were transfected with the spike expression plasmid and subsequently infected with rVSV-*firefly*-luciferase (Kerafast, EH1020-PM) as previously described[1,2]. The pseudovirus-containing culture supernatant was harvested and centrifuged at 500 g for 5 min to remove cell debris. The cell-free supernatant was then filtered through a 0.45 μm polyethersulfone membrane and stored at -150°C in 100 μL aliquots for future use.

For the neutralization assay, Hela/ACE2-11 cells (Gift from Dr. Guangxiang Luo, University of Alabama) were seeded in 96-well plates in culture medium and allowed to reach approximately 85% confluence before use in the assay (24 h later). Sera were 3-fold serially diluted in medium (dilutions ranged from 1:50 to 1:36,450) and incubated with VSV based-pseudovirus at 37 °C for 30 minutes. The virus-sera mix was subsequently used to infect Hela/ACE2-11 cells for 36 h at 37 °C after which cells were washed with PBS and lysed with Passive Lysis Buffer (Promega). Firefly luciferase activity (relative luminescence unit; RLU) in lysates was measured using the Luciferase Assay System (Promega) with EnVision Multimode Plate Reader (PerkinElmer). The obtained RLU was normalized to those derived from cells infected with pseudovirus only. The half-maximal inhibitory dilution for serum ($ID_{50}$) was determined using four-parameter logistic curve (GraphPad Prism Version 9.1.1).

#### SARS-CoV-2 Spike Human IgG ELISA

The SARS-CoV-2 Spike S1 human IgG from vaccinated human sera was quantified with ELISA following the protocol of the manufacturer. Briefly, human sera were diluted at 1:5000 and 1:10000 with assay buffer, then added to SARS-CoV-2 S1 pre-coated wells and incubated at room temperature for 2 h, followed by incubation with biotinylated SARS-CoV-2 S1 detection antibody for 1 h and Avidin-HRP for 30 min. Peroxidase substrate solution (TMB) and 1M $H_2SO_4$ stop solution were used and the absorbance (OD 450 nm and 570 nm) was read by a microplate reader (Spectra Max i3, Molecular Devices, Sunnyvale, CA, USA). Lyophilized SARS-CoV-2 Spike S1 human IgG was 2-fold serially diluted starting from 15 ng/mL and used as standard for accurate quantitation of SARS-CoV-2 Spike S1 human IgG.

**Statistical Analysis**

GraphPad Prism software (Version 9.1.1) was used for statistical analysis. Wilcoxon matched-pairs signed rank test was used for two-group analysis. Pearson's correlation coefficients were calculated. *P* values less than 0.05 were statistically significant.

**Figure S1**

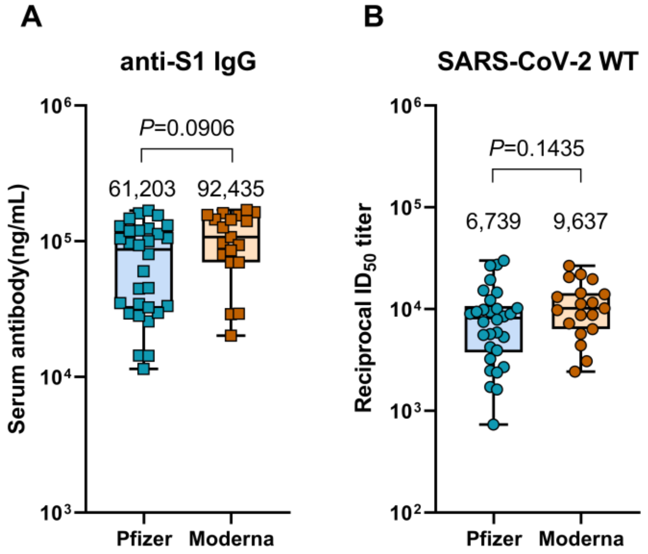

**Figure S1**. **Specific anti-SARS-CoV-2 S1 IgG elicited by Pfizer and Moderna vaccine and neutralizing activity against SARS-CoV-2 WT.**

(**A**) Shown is specific anti-SARS-CoV-2 S1 IgG in sera collected from Pfizer (N=30) and Moderna (N=19) vaccinated subjects approximately 3 weeks to 2 months after second dose of vaccination. The numbers over the dot of each group are the geometric mean. (**B**)Shown is 50% pseudovirus neutralizing titer (50% inhibitory dilution, $ID_{50}$) against recombinant VSV–based SARS-CoV-2 pseudovirus bearing the Wuhan-1 (wild-type, WT) spike protein in sera collected from Pfizer and Moderna vaccinated subjects. The numbers over the dot of each group are the geometric mean titers (GMTs). Box plots indicate the median and interquartile range (IQR); the whiskers represent 1.5 times the IQR.

**Figure S2**

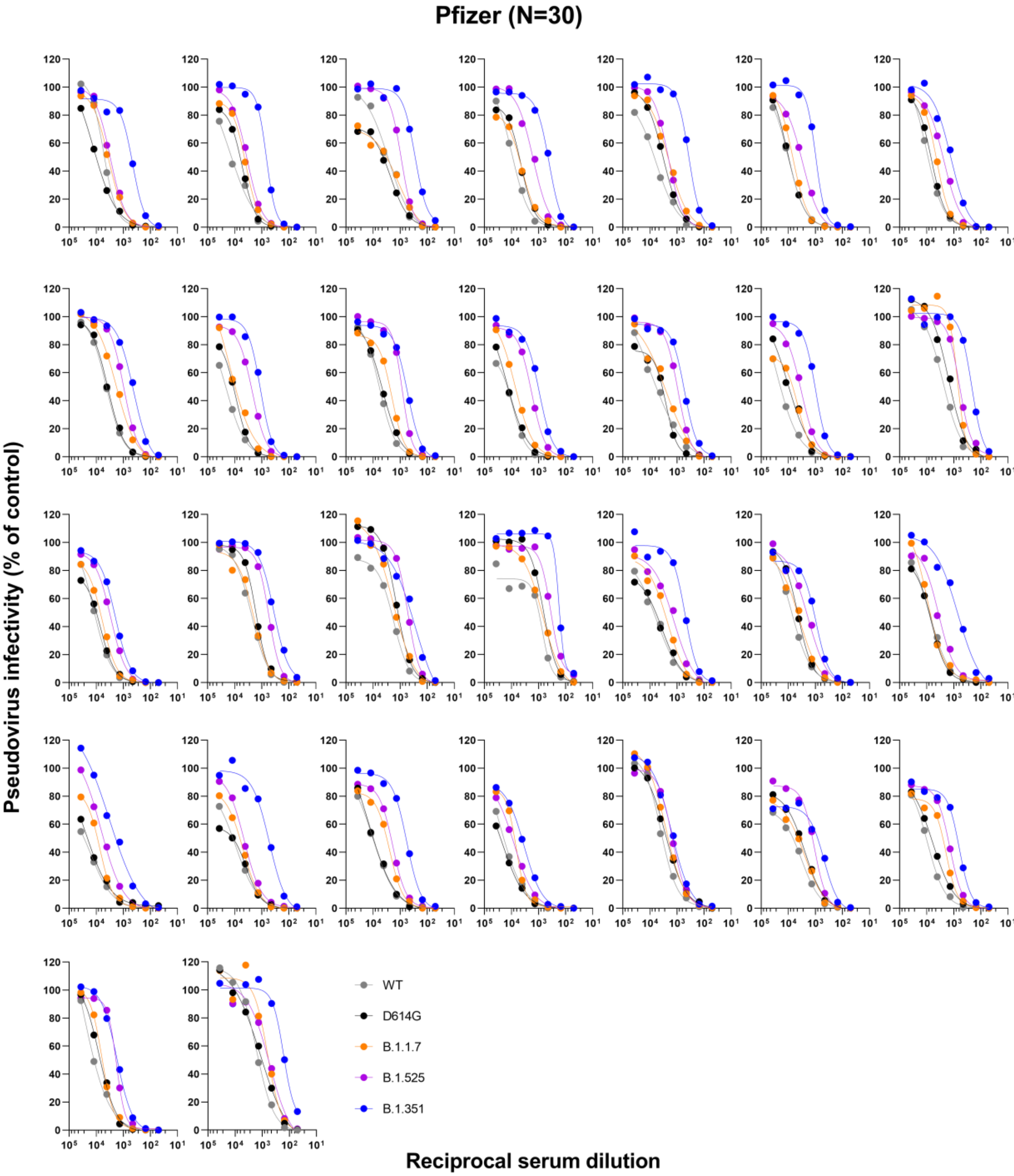

Pfizer (N=30)
Pseudovirus infectivity (% of control)
Reciprocal serum dilution
WT
D614G
B.1.1.7
B.1.525
B.1.351

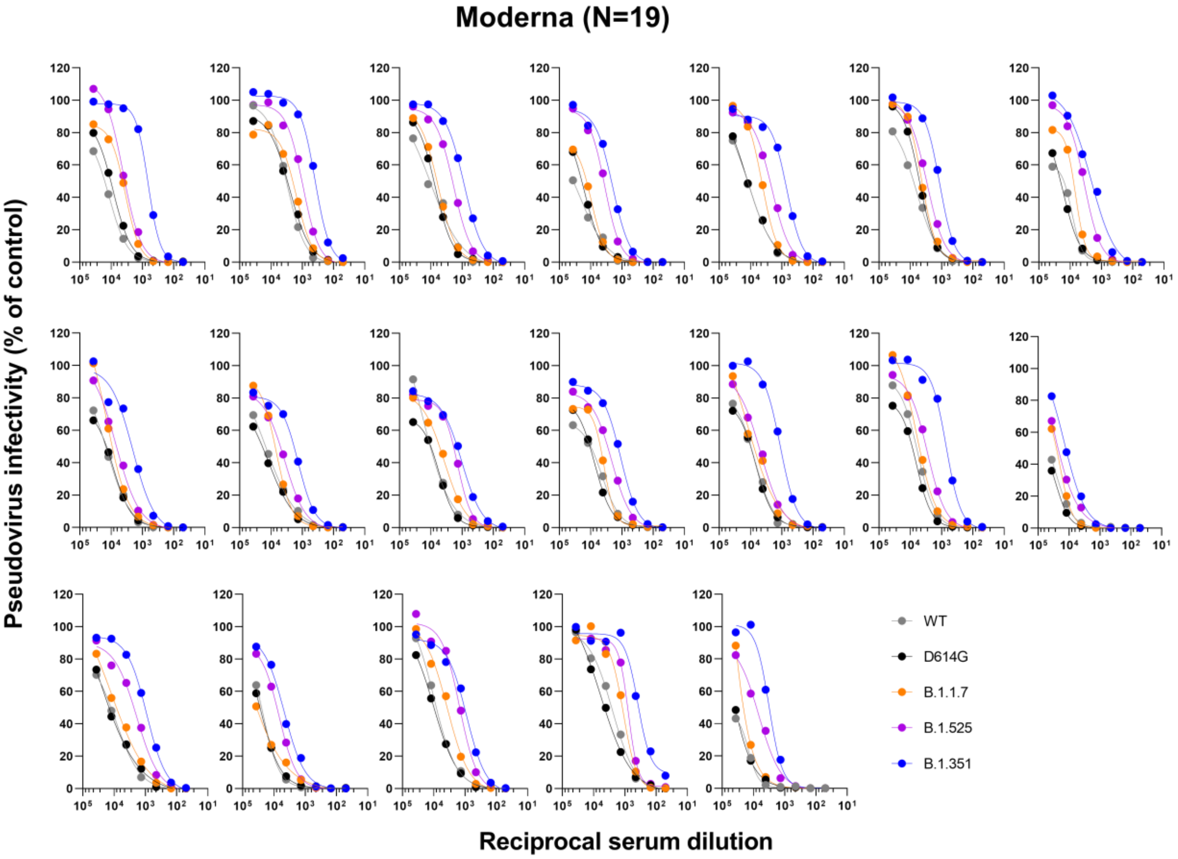

**Figure S2. Neutralization curves of human sera in the VSV-based pseudovirus neutralization assay.** Sera were collected from Pfizer (N=30) and Moderna (N=19) vaccinated subjects approximately 3 weeks to 2 months after second dose of vaccination. Neutralization was measured against recombinant VSV-based pseudovirus bearing the full-length spike protein of SARS-CoV-2 WT, D614G, B.1.1.7, B.1.525, or B.1.351 variant. Each graph represents an individual participant, as indicated. Each data point is an average from duplicate wells. The obtained RLU were normalized to those derived from cells infected with pseudovirus only (control), and subsequently analyzed using four-parameter logistic curve (GraphPad Prism Version 9.1.1).

### Figure S3

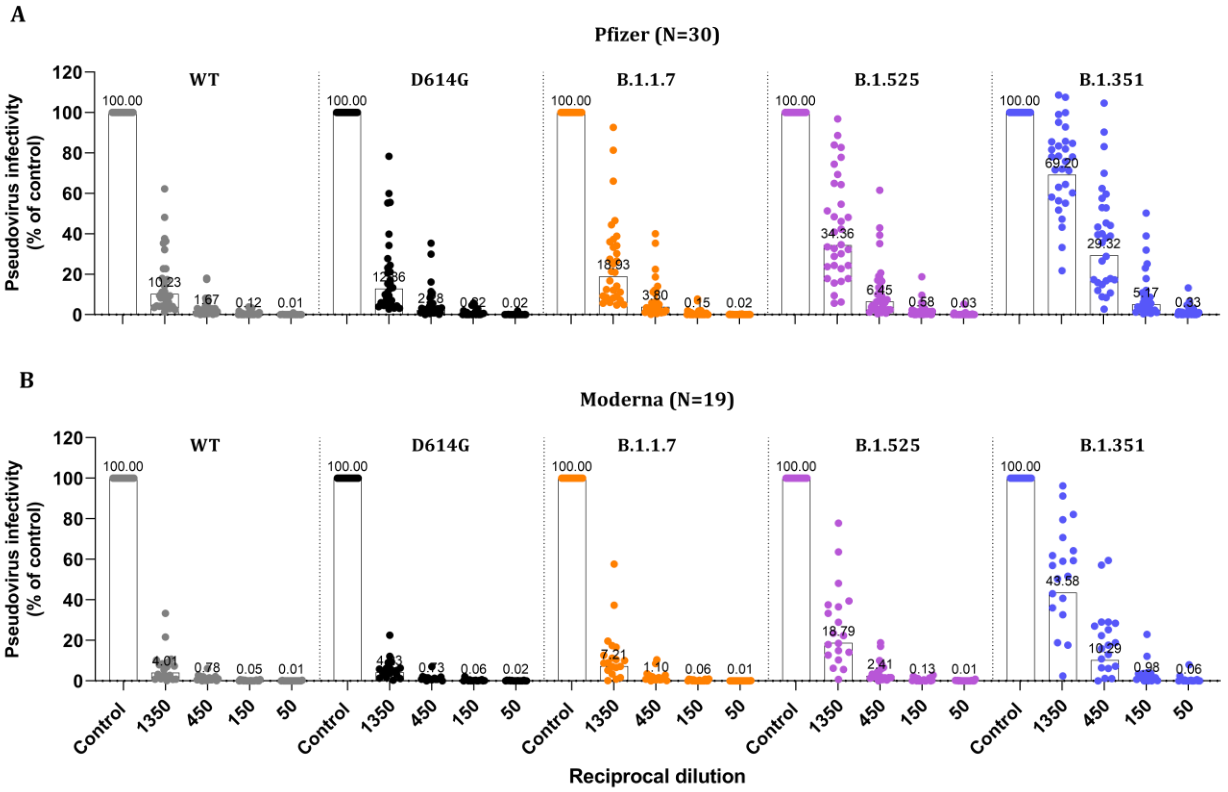

**Figure S3. Neutralizing activity of diluted sera against SARS-CoV-2 WT and its variants.**

Sera were collected from Pfizer (N=30) and Moderna (N=19) vaccinated subjects, and 3-fold serially diluted in medium starting with a 1:50 dilution. Neutralization was measured against recombinant VSV-based pseudovirus bearing the full-length spike protein of SARS-CoV-2 WT, D614G, B.1.1.7, B.1.525, or B.1.351 variant. Firefly luciferase activity (relative luminescence unit; RLU) in lysates was measured using the Luciferase Assay System (Promega) with EnVision Multimode Plate Reader (PerkinElmer). The obtained RLU was normalized to those derived from cells infected with pseudovirus only (control). Each dot represents an individual participant. The numbers over the dot of each group are the geometric mean of the relative infectivity.

**Table S1**

**Characteristics of study subjects**

| Characteristics | Pfizer vaccinee | Moderna vaccinee |
|---|---|---|
| **No. of subjects** | 30 | 19 |
| **Age (median, range)** | 36.0 (21.0-73.0) | 39.0 (20.0-65.0) |
| **Sex** | | |
| Male (%) | 11 (36.7) | 7 (36.8) |
| Female (%) | 19 (63.3) | 12 (63.2) |
| **Days post 2nd Dose (Median day, range)** | 30.5 (22-68) | 35.0 (24.0-49.0) |
| **Race** | White: 16<br>Black or African American: 2<br>Asian: 12 | White: 11<br>Black or African American: 1<br>Asian: 7 |

**Table S2**

**Spike mutations in SARS-CoV-2 variants**

| SARS-CoV-2 | Amino Acid Changes in Spike |
|---|---|
| **Wild Type (WT)**<br>(Wuhan-1 reference strain) | |
| **B.1** | D614G |
| **B.1.1.7**<br>(20I/501Y.V1, VOC 202012/01) | ΔHV69-70-ΔY144-N501Y-A570D-D614G-P681H-T716I-S982A-D1118H |
| **B.1.351**<br>(20H/501Y.V2) | L18F-D80A-D215G-ΔLAL242-244-R246I-K417N-E484K-N501Y-D614G-A701V |
| **B.1.525**<br>(20C) | Q52R-ΔHV69-70-ΔY144-E484K-D614G-Q677H-F888L |
